# Development of LRRK2 designed ankyrin-repeat proteins

**DOI:** 10.1101/2023.11.14.567123

**Authors:** Verena Dederer, Marta Sanz Murillo, Eva P Karasmanis, Kathryn S Hatch, Deep Chatterjee, Franziska Preuss, Kamal R Abdul Azeez, Landon Vu Nguyen, Christian Galicia, Birgit Dreier, Andreas Plückthun, Wim Versees, Sebastian Mathea, Andres E Leschziner, Samara L Reck-Peterson, Stefan Knapp

## Abstract

Leucine rich repeat kinase 2 (LRRK2) is a large multidomain protein containing two catalytic domains, a kinase and a GTPase, as well protein interactions domains, including a WD40 domain. The association of increased LRRK2 kinase activity with both the familial and sporadic forms of Parkinson’s disease (PD) has led to intense interest in determining its cellular function. However, small molecule probes that can bind to LRRK2 and report on or affect its activity are needed. Here, we identified a series of high-affinity LRRK2-binding designed ankyrin-repeat proteins (DARPins). One of these DARPins (E11) bound to the LRRK2 WD40 domain with high affinity. LRRK2 bound to DARPin E11 showed improved behavior on cryo-EM grids, resulting in higher resolution LRRK2 structures. DARPin E11 did not affect the catalytic activity of a truncated form of LRRK2 in vitro but decreased the phosphorylation of Rab8A, a LRRK2 substrate, in cells. We also found that DARPin E11 disrupts the formation of microtubule-associated LRRK2 filaments in cells, which are known to require WD40-based dimerization. Thus, DARPin E11 is a new tool to explore the function and dysfunction of LRRK2 and guide the development of LRRK2 kinase inhibitors that target the WD40 domain instead of the kinase.

## Introduction

Mutations in Leucine-Rich Repeat Kinase 2 (*LRRK2*) are one of the most common causes of familial Parkinson’s disease (Paisán-Ruíz et al., 2004; Zimprich et al., 2004). LRRK2 is a large multi-domain protein. Its N-terminal half contains armadillo, ankyrin, and leucine-rich repeats, while its C-terminal catalytic half contains GTPase (<u>R</u>as <u>O</u>f <u>C</u>omplex, or ROC), <u>C</u>-terminal <u>O</u>f <u>R</u>OC (COR-A and COR-B), kinase, and WD40 domains (**Figure 1A**). LRRK2 forms dimers at two interfaces: between its WD40 domains and between its COR-B domains (Deniston et al., 2020; Myasnikov et al., 2021; Zhang et al., 2019); these interfaces are important for LRRK2 to form filaments on microtubules, which is enhanced by high expression levels of LRRK2, in the presence of LRRK2 type-I kinase inhibitors, and by most PD variants of LRRK2 (Blanca Ramírez et al., 2017; Deniston et al., 2020; Kett et al., 2012; Watanabe et al., 2020). Most PD-linked variants of LRRK2 also increase its kinase activity (Gloeckner et al., 2005; Sheng et al., 2012; Steger et al., 2016; West et al., 2005). Thus, small molecules that inhibit LRRK2’s kinase activity are promising drug candidates for PD treatment (Taymans et al., 2023).

**Figure 1.**
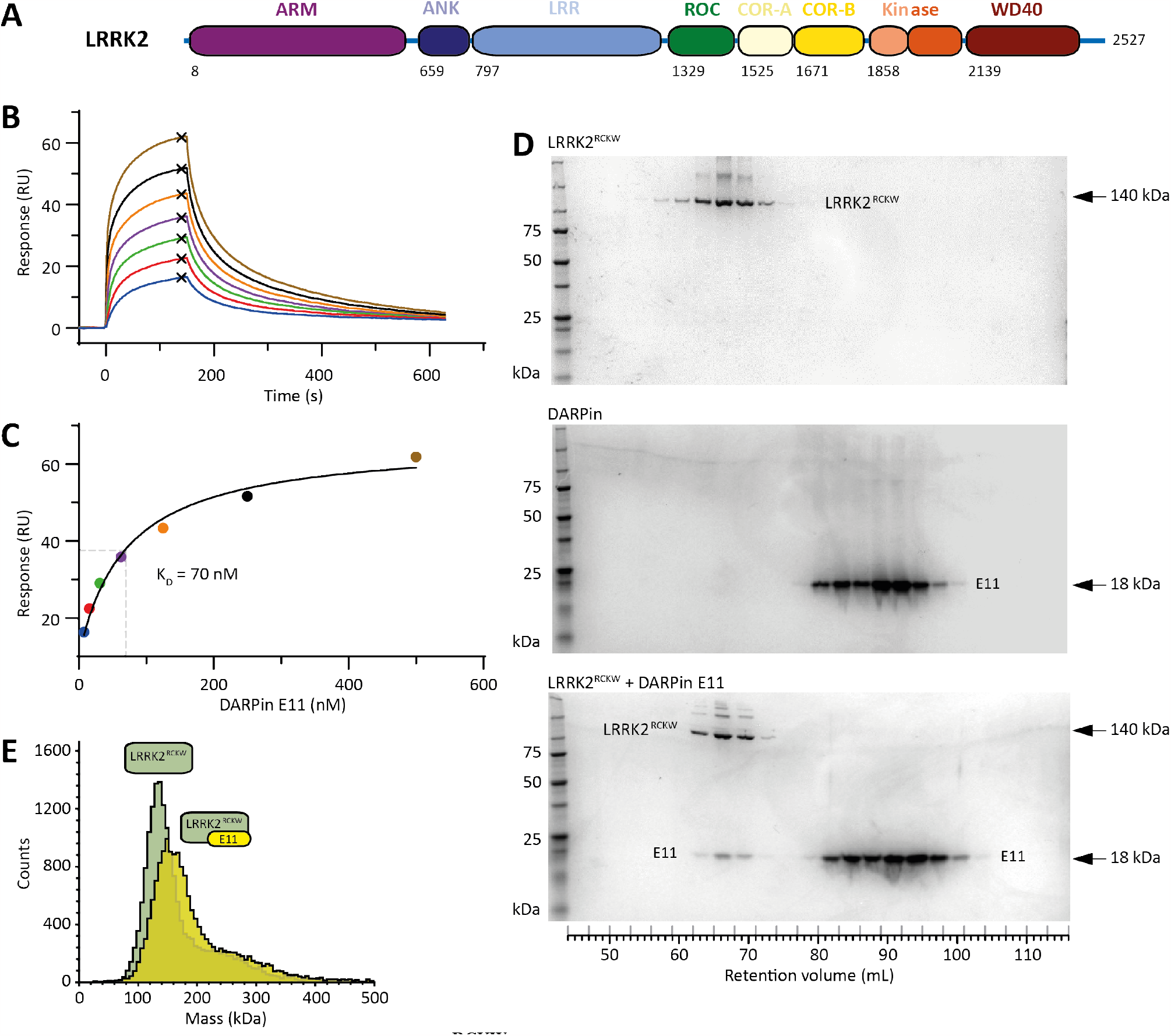
DARPin E11 binds to LRRK2^RCKW^ with high affinity. **A**. Domain architecture of the LRRK2 protein. The same color scheme is used in all figures. The first residue of each domain is indicated. **B, C**. Surface plasmon resonance analysis of DARPin E11 binding to LRRK2^RCKW^. Recombinant LRRK2^RCKW^ was immobilized on a sensor chip and varying concentrations of DARPin E11 were added in the mobile phase. Sensograms are shown (B) and the plateau values from the doseresponse curves (C) are shown and fit to the Langmuir equation. A dissociation constant (K_D_) of 70 nM was determined. **D**. The LRRK2^RCKW^:E11 complex was subjected to size exclusion chromatography (SEC). Elution fractions were analyzed by SDS PAGE and visualized by Coomassie staining. The co-elution of LRRK2^RCKW^ and DARPin E11 showed that the complexes were stable during the SEC run. Molecular weight markers are noted on the left and the molecular weights of LRRK2^RCKW^ and DARPin E11 on the right. **E**. Analysis of the LRRK2^RCKW^:E11 complex by mass photometry. The molecular weight obtained from the yellow peak corresponds to that of a 1:1 LRRK2^RCKW^:E11 complex.

Given the complexity of LRRK2 regulation and signaling, it is necessary to study these processes in cellular environments. Small molecule and protein-based binders have been developed for this purpose. The most important of these in terms of therapeutic development are the type-I kinase inhibitors that bind to the active-like state of LRRK2. Several type-I inhibitors have been developed, including MLi-2 (Fell et al., 2015) and DNL201 (developed as GNE0877) (Estrada et al., 2014). DNL151 (renamed BIIB122), which is related to DNL201, has entered phase 2b clinical trials (clinicaltrials.gov). In contrast, there are currently no published LRRK2-selective type-II inhibitors that target the LRRK2 inactive state. An example of a high-affinity, but not selective type-II inhibitor for LRRK2 is rebastinib (Schmidt et al., 2021). In addition to kinase inhibitors, other LRRK2 tools have been developed. These include a proteolysis targeting chimera (PROTAC) called XL01126 (Liu et al., 2022), a small molecule that binds to its ROC GTPase domain called FX2149 (Li et al., 2015), and nanobodies (Chaikuad et al., 2014; Singh et al., 2022).

In addition to nanobodies, designed ankyrin repeat proteins (DARPins) are a class of protein-based binders (Plückthun, 2015). DARPins are inspired by the naturally occurring ankyrin repeat (AR) domains. They comprise two to three ARs that each contain randomized regions (7 randomized residues per repeat), flanked by capping repeats with a hydrophilic surface, and newer versions of the libraries also contain randomized loops and randomized regions in the capping repeats (Plückthun, 2015; Schilling et al., 2014). The theoretical diversity far exceeds the amount of DNA encoding it, but with ribosome display (Dreier and Plückthun, 2012; Plückthun, 2012), libraries are limited by the number of ribosomes in the reaction mix (typically 10^12^) used for selecting specific binders (Plückthun, 2015). Binders to several hundred different targets have been reported. As specific binding proteins, DARPins have similar applications as antibodies but feature several advantages: they are smaller in size, are very stable and rigid, can be produced in *E. coli*, harbour no disulfide bonds, and can thus be used in the reducing cytoplasm and are easier to engineer to many different formats. Some DARPins have entered clinical trials (Kunimoto et al., 2020; Radom et al., 2019) and they have been used as conformation-specific sensors in living cells (Kummer et al., 2013; Strubel et al., 2022).

DARPins have also been used to modulate sample properties in cryo-EM applications. DARPins can be designed to selectively bind to and stabilize specific conformations or states of the target molecules (Strubel et al., 2022). They have been useful for studying conformational changes or for understanding the molecular determinants of protein activation and autoinhibition (Qi et al., 2022). Currently, proteins smaller than about 50 kDa present a challenge for structural elucidation by cryo-EM. Recent reports have shown that small proteins can be visualized attached to a large, symmetric base or cage via a DARPin (Liu et al., 2018; Vulovic et al., 2021; Yao et al., 2019). The additional molecular mass and features contributed by the DARPin can also improve the signal-to-noise ratio and facilitate the determination of high-resolution structures. Finally, the adoption of a limited number of orientations on the grid by a target molecule, resulting from the interaction with the air-water interface, remains the most common and significant challenge in cryo-EM. DARPins could help increase the number of orientations adopted by a target molecule by changing its surface properties.

Here, we describe the identification and characterization of DARPins that bind to LRRK2. We report the characterization of DARPin E11, which showed one of the highest affinities for LRRK2. DARPin E11 did not affect the kinase activity of a truncated LRRK2 construct in vitro. Surprisingly, however, it inhibited full length LRRK2’s ability to phosphorylate Rab8A, one of LRRK2’s physiological substrates, when expressed in cells. Expressing DARPin E11 in cells also prevented the formation of microtubule-associated LRRK2 filaments, which are mediated by a WD40-WD40 interaction that overlaps with the E11 binding site on the WD40 domain. Our cryo-EM analysis of the LRRK2:E11 complex indicated that addition of the DARPin increased the number of orientations adopted by LRRK2 on cryo-EM grids, leading to an improved cryo-EM structure. Thus, DARPin E11 is a new tool with potential applications in structural studies of LRRK2:ligand complexes, functional studies of LRRK2 in cells, and the design of LRRK2 inhibitors that target its WD40 domain.

## Methods

### Production of the biotinylated LRRK2^RCKW^ protein

The DNA coding for the LRRK2 residues 1327 to 2527 (OHu107800 from Genscript) was PCR-amplified using the forward primer TACTTCCAATCCATGAAAAAGGCTGTGCCTTATAACCGA and the reverse primer TATCCACCTTTACTGCTCTCAACAGATGTTCGTCTCATTTTTTCA. The T4 polymerase-treated amplicon was inserted into the transfer vector pFB-Bio5 (SGC) by ligation-independent cloning. The resulting plasmid was utilized for the generation of recombinant Baculoviruses according to the Bac-to-Bac expression system protocol (Invitrogen). Exponentially growing Sf9 cells (2 x 10^6^ cells/mL in Lonza Insect-XPRESS medium supplemented with 0.1 mM biotin) were infected with a high-titer Baculovirus suspension. After 66 hours of incubation (27°C and 90 rpm), cells were harvested by centrifugation. The expressed protein construct contained an N-terminal His_6_-tag, cleavable with TEV protease. For LRRK2^RCKW^ purification, the pelleted Sf9 cells were washed with PBS, re-suspended in lysis buffer (50 mM HEPES pH 7.4, 500 mM NaCl, 20 mM imidazole, 0.5 mM TCEP, 5% glycerol) and lysed by sonication. The lysate was cleared by centrifugation and loaded onto a Ni NTA column. After vigorous rinsing with lysis buffer the His_6_-tagged protein was eluted in lysis buffer containing 300 mM imidazole. The eluate was treated with TEV protease to cleave the His_6_-tag and dialysed overnight in storage buffer (20 mM HEPES pH 7.4, 150 mM NaCl, 0.5 mM TCEP, 5% glycerol) to reduce the imidazole concentration. Contaminating proteins, uncleaved LRRK2^RCKW^ protein and TEV protease were removed with another Ni NTA step. Finally, LRRK2^RCKW^ was concentrated and subjected to gel filtration in storage buffer using an AKTA Xpress system combined with an S200 gel filtration column. The elution volume 69.3 mL indicated the protein to be monomeric in solution. The final yield as calculated from UV absorbance was 0.6 mg biotinylated LRRK2^RCKW^/L insect cell medium.

### Selection of DARPins

The DARPin library consisted of N2C and N3C DARPins with randomized loops and randomized capping repeats (Plückthun, 2015). The number of ribosomes used in ribosome display allow one to select from about 10^12^ members. To enrich LRRK2-binding DARPins, the biotinylated LRRK2^RCKW^ protein was immobilized to MyOne T1 streptavidin-coated beads (Thermo Fisher Scientific). The ribosome display selections were performed using a KingFisher Flex MTP96 well platform as described previously (Dreier and Plückthun, 2012; Strubel et al., 2022). Four rounds of selection were performed with decreasing amounts of LRRK2^RCKW^. In the third round, low off-rate DARPins were enriched by adding non-biotinylated LRRK2^RCKW^ as a competitor. In the fourth round, less stringent washing conditions than in the other rounds were applied to recover binders (Dreier and Plückthun, 2012; Strubel et al., 2022). The resulting pool of DARPin cDNAs was cloned into the vector pQE30 (Qiagen) allowing for the expression of DARPins with an N-terminal His-tag and a C-terminal FLAG-tag. *E. coli* XL1-Blue was transformed with this enriched pool, and 380 single colonies were isolated. To confirm DARPin binding to LRRK2^RCKW^, a homogeneous time-resolved fluorescence (HTRF) assay was performed according to a previously established protocol (Dreier and Plückthun, 2012; Strubel et al., 2022). In brief, the 380 DARPins were expressed in 1-mL scale in deep well plates, cells were harvested by centrifugation, and lysed in a lysozyme- and detergent-containing buffer. The lysates were cleared by centrifugation. Then, biotinylated LRRK2^RCKW^ was added to allow for the formation of LRRK2^RCKW^:DARPin complexes. After mixing with the FRET donor Streptavidin-Tb cryptate (610SATLB, Cisbio) and the FRET acceptor mAb anti-FLAG M2-d2 (61FG2DLB, Cisbio) and an incubation time of 30 minutes, the samples were analyzed using a Varioskan LUX Multimode Microplate reader (Thermo Scientific). The HTRF ratios were obtained by dividing the acceptor fluorescence signal by the donor fluorescence signal. The 20 DARPins with the highest HTRF ratios were chosen, sequenced (**Table S1**) and expression scaled up for further analysis.

### Determination of DARPin affinities by surface plasmon resonance (SPR)

The SPR analysis was performed on a Biacore T200 (Cytiva Life Sciences). 850 RU of biotinylated LRRK2^RCKW^ was loaded onto a Series S CM5 chip coated with Streptavidin. The chip was equilibrated with running buffer containing 20 mM HEPES pH7.4, 150 mM NaCl, 0.5 mM TCEP, 20 μM GDP and 2.5 mM MgCl_2_. A titration was performed for DARPin E11. The DARPin was allowed to bind over the surface at a flow rate of 30 μL/min for 150 or 180 seconds and disassociation was followed for 480 seconds. The sensograms were double-reference subtracted and analyzed by a 1+1 kinetic model and a 1+1 Langmuir adsorption isotherm model for DARPin E11.

### Determination of DARPin affinities by bio-layer interferometry (BLI)

The measurements were performed using an Octet Red96 (FortéBio) system at 25°C in a buffered solution containing 50 mM HEPES pH 8.0, 150 mM NaCl, 10 mM MgCl_2_, 5% glycerol, 0.1% bovine serum albumin (BSA), 0.05% Tween 20 and 500 mM GDP. Binding of DARPins to LRRK2^RCKW^ was measured using biotinylated LRRK2^RCKW^ protein loaded onto streptavidin-coated (SA) biosensors. The protein was immobilized to the sensors from a fixed LRRK2^RCKW^ concentration of 5 μg/mL, and binding curves were measured by varying concentrations of DARPin E11 from 15 to 1000 nM. The association/dissociation traces were fitted using the global option (implemented in the FortéBio Analysis Software). The resulting R_eq_ values were subsequently plotted against the DARPin concentration and used to derive the K_D_ values from the corresponding dose–response curves by fitting on a Langmuir model. Dose-response curves were measured in triplicates. Final figures were generated using GraphPad Prism7.

### Mass photometry analysis

All mass photometry data was acquired on a Refeyn Two MP mass photometer with the Refeyn Acquire MP Software 2023.1.1. The measurements were carried out in reusable gaskets (Grace Bio Labs, CW-50R-1.0 50-3DIA x1 mm) on high precision cover slips which had been cleaned with isopropanol, 70% ethanol and Milli-Q water. 20 μL buffer (20 mM HEPES pH 7.4, 150 mM NaCl, 2.5 mM MgCl_2_, 5% glycerol, 20 μM GDP and 0.5 mM TCEP) was added to the gaskets. After focusing, 10 μL of the buffer drop was exchanged for the protein solution to be measured. For landing events, 2997 image frames were acquired during 60 s measurement time at 499.5 Hz. All data were analyzed with the RefeynDiscover MP Software 2023.1.2. Ratiometric contrast values were converted into molecular mass using a standard mass calibration (using BSA, CA and IgG as reference).

### Purification of LRRK2^RCKW^ for Cryo-EM

Sf9 cells expressing His_6_-Z-TEV-LRRK2^RCKW^ were washed with PBS, resuspended in lysis buffer (50 mM HEPES pH 7.4, 500 mM NaCl, 20 mM imidazole, 0.5 mM TCEP, 5% glycerol, 5 mM MgCl_2_, 20 μM GDP) and lysed by homogenization. The supernatant was cleared by centrifugation and loaded onto a Ni-NTA (Qiagen) column. After rinsing with lysis buffer, the His_6_-Z-tagged protein was eluted in lysis buffer containing 300 mM imidazole. The eluate was then diluted to 250 mM NaCl with a dilution buffer (50mM HEPES pH 7.4, 0.5mM TCEP, 5% glycerol, 5mM MgCl_2_, 20μM GDP) and loaded onto an SP sepharose column. His_6_-Z-TEV-LRRK2^RCKW^ was eluted with a gradient from 250 mM to 2.5 M NaCl in dilution buffer and treated with TEV protease overnight to cleave the His_6_-Z-tag. Contaminating proteins, the cleaved tag, uncleaved protein and TEV protease were removed by another combined SP sepharose Ni-NTA step. Finally, LRRK2^RCKW^ was concentrated and subjected to gel filtration in 20 mM HEPES pH 7.4, 700 mM NaCl, 0.5 mM TCEP, 5% glycerol, 2.5 mM MgCl_2_, 20 μM GDP using an AKTA Xpress system combined with an S200 gel filtration column. The final yield, as calculated from UV absorbance, was 2.1 mg of LRRK2^RCKW^ per liter of insect cell medium.

### Preparation of LRRK2^RCKW^:E11 complex for structural studies

LRRK2^RCKW^ was exchanged into buffer A (20 mM HEPES pH 7.4, 150 mM NaCl, 2.5 mM MgCl_2_, 5% glycerol, 20 μM GDP and 0.5 mM TCEP). DARPin E11 was incubated with LRRK2^RCKW^ in a ratio of 1:1.25 for 10 minutes at RT and then placed at 4°C and incubated for 15 minutes. After that, the sample was diluted with cold buffer A to a final LRRK2^RCKW^ concentration of 6 μM. The sample was used to make cryo-EM grids immediately after dilution.

### Cryo-EM sample preparation

3.5 μL of LRRK2^RCKW^:E11 complex was applied to glow-discharged UltraFoil 300 mesh R1.2/1.3 grids (Quantifoil) and incubated in a FEI Vitrobot IV at 4°C and 95% humidity for 20 sec. The grids were blotted for 4 s at an offset of -20 mm on both sides and vitrified by plunging into liquid ethane cooled down to liquid nitrogen temperature. Data were collected on an FEI Talos Arctica operated at 200 kV and equipped with a K2 summit direct electron detector (Gatan), using Leginon (Suloway et al., 2005). Images were acquired at defocus values varying between -1.0 and -2.0 μm at a nominal magnification of 36000x, yielding a pixel size of 1.16 Å. The camera was operated in dose-fractionation counting mode collecting 40 frames per movie, with a total dose of 60 e^-^ per Å^2^.

### Cryo-EM data processing

2,468 movies were collected and aligned using MotionCor2 (Zheng et al., 2017) and CTF parameters were estimated using Ctffind4 (Rohou and Grigorieff, 2015). Micrographs with a CTF estimation worse than 5 Å were discarded. Particles obtained with blob picker in cryosparc (Punjani et al., 2017) were used to train a Topaz model (Bepler et al., 2019). After inspection, particle extraction and several rounds of 2D classification, approximately 280,000 particles were extracted with a 288-pixel box and used for subsequent processing. After ab initio and heterogeneous refinement to separate particles with and without DARPin E11, 185,601 particles were selected and used as an input for a Non-Uniform Refinement. A final local refinement using a mask for the C-lobe of the kinase, the WD40 domain and E11 DARPin yielded a map at 3.6 Å.

### Model building

The available structure of LRRK2^RCKW^ (PDB ID 6VP7) was split into domains and fitted into the 3D maps using UCSF ChimeraX (Pettersen et al., 2021). For DARPin E11, we generated an initial model using AlphaFold (Jumper et al., 2021; Mirdita et al., 2021) and used it as starting point for model building, which was performed in Coot (Emsley et al., 2010). The built structures were refined using real-space refinement in Phenix (Liebschner et al., 2019). Refinement statistics are summarized in **Table S2**. Figures were prepared using USCF ChimeraX (Pettersen et al., 2021).

### Kinase activity assay

To investigate the impact of DARPin binding on LRRK2 kinase activity, a mass spectrometry (MS)-based activity assay was performed. Phosphorylation of the physiological LRRK2 substrate Rab8A was measured. The reaction was set up as follows: 5 μM recombinant Rab8A was incubated with 50 nM LRRK2^RCKW^ and a concentration series of the respective DARPin ranging from 0.016 to 1 μM in reaction buffer (20 mM HEPES pH 7.4, 100 mM NaCl, 0.5 mM TCEP, 0.5 % glycerol, 20 μM GDP, 2.5 mM MgCl_2_). The phosphorylation reaction was started by adding ATP to the final concentration 1 mM. After incubation at room temperature for 3.5 h the reaction was stopped by adding an equal volume of MS buffer (dH2O with 0.1 % formic acid). The Rab8A phosphorylation grade was analyzed by MS with an Agilent 6230 Electrospray Ionization Time-of-Flight mass spectrometer coupled with the liquid chromatography unit 1260 Infinity. The sample was applied via a C3 Column and eluted at 0.4 mL/min flow rate using a solvent gradient of water to acetonitrile with 0.1% formic acid. Data were acquired with the MassHunter LC/MS Data Acquisition software and analyzed with the BioConfirm vB.08.00 tool (both Agilent Technology).

### Immunofluorescence, confocal microscopy, and image analysis

293T cells were plated on fibronectin-coated glass 22 mm x 22 mm coverslips and grown for 24 hours before transfection with PEI and 800 ng of GFP-LRRK2 alone or co-transfected with 400 ng 8xHis-DARPinE11-3xFLAG (in pCDNA3.1). After 24–48 hours, the cells were incubated at 37 °C with DMSO or MLi-2 (Tocris; 2 μM) for 2 hours. The cells were fixed with prewarmed 3% PFA, 4% sucrose, in 1X PBS for 10 minutes at room temperature. The coverslips were then rinsed twice and washed twice with 1X PBS and quenched with 0.4% NH_4_Cl for 10 minutes. After washing with PBS, the cells were incubated with blocking and permeabilizing buffer (2% BSA, 0.1% Triton X-100 in 1X PBS) for 20 minutes at room temperature. A primary Rabbit anti-FLAG polyclonal antibody (ptg labs 20543-1-AP) was diluted 1:200 in antibody dilution buffer (2% BSA in 1x PBS) and incubated with the cells for 3 hours at room temperature. Following primary antibody incubation, the coverslips were washed three times with 2% BSA in PBS 1X and incubated with 1:500 goat anti-rabbit Alexa568 secondary antibody (diluted in antibody dilution buffer) for 1 hour at room temperature. After secondary antibody incubation, the cells were treated with 1:1000 DAPI in PBS for 10 minutes and then washed five times with 1X PBS before mounting on glass slides using FluorSave (EMD Millipore Sigma). The coverslips were left to dry at least one hour and imaged immediately or stored at 4ºC for later imaging. Mounted cover slips were blinded before imaging to prevent bias in image acquisition. Imaging of blinded samples was done using a Yokogawa W1 confocal scanhead mounted to a Nikon Ti2 microscope with an Apo ×60 1.49-NA objective. The microscope was run with NIS Elements using the 488 nm, 561nm and 405 nm lines of a six-line (405 nm, 445 nm, 488 nm, 515 nm, 561 nm, and 640 nm) LUN-F-XL laser engine and a Prime95B camera (Photometrics). 16-20 areas were imaged per sample. Cells expressing both LRRK2 (positive for GFP fluorescence signal) and DARPin E11 (stained with anti-FLAG antibody) were assessed for the presence or absence of LRRK2 filaments in ImageJ2 (version 2.14.0). Maximum projections were generated from confocal z-stacks as a guide. The number of cells with filaments was divided by the total number of cells co-expressing LRRK2 and DARPin E11 to determine the percent of cells with filaments per replicate. Three or four replicates were obtained for each condition on at least two separate days. The slides were imaged blinded and analyzed by two different researchers. Unblinding took place only after all analyses were completed. Data were plotted and statistics were generated in GraphPad Prism (version 10.0.3).

### LRRK2 kinase assay in cells

293T cells were transfected with 1000 ng of wild type LRRK2 and 500 ng GFP-Rab8A or 1000 ng LRRK2, 500 ng GFP-Rab8A and 500 ng of 8xHis-DARPin E11-3xFLAG using PEI. After 48 hours, cells transfected only with LRRK2 and GFP-Rab8A (without DARPin E11) were incubated at 37ºC, 5% CO2 with either DMSO or MLi-2 (Tocris; 2 μM) for 1 hour. Cells were washed twice with room temperature PBS one time and a final time with cold PBS. Cells were lysed in cold RIPA lysis buffer (50 mM Tris pH 8.0, 150 mM NaCl, 1 % Triton X-100, 0.1% SDS, 0.5% Sodium Deoxycholate, 1mM DTT) in the presence of cOmplete mini EASYpack protease inhibitor cocktail and the PhosSTOP EASYpack phosphatase inhibitor cocktail (Roche). Resuspended cells were vortexed for 30 seconds six times and incubated on a rotator for 15 minutes at 4ºC. Cell lysates were then spun down at 13,000 x g for 15 minutes at 4ºC. SDS PAGE loading buffer was added to the supernatant and was either used immediately or stored at -20ºC. 15 μl of sample were loaded into a NuPAGE 4-12% gradient Bis-Tris gel (Invitrogen). Protein was transferred onto PVDF transfer membrane (0.45 μm pore size) for 100 minutes at 200 mA at 4ºC. The membrane was cut directly above the 70 kDa molecular weight marker band to separate LRRK2 from the rest of the membrane. The upper portion of the membrane containing LRRK2 was blocked with 5% BSA for 1 hour, at room temperature and incubated with 1:1000 rabbit anti-LRRK2 monoclonal antibody (Abcam c41-2) diluted in 1% BSA O/N at 4°C. The lower part of the membrane (below 70 kDa) was blocked in 5% milk for one hour at room temperature and incubated with a monoclonal mouse anti-GFP antibody (Santa Cruz Biotechnology sc-9406, 1:2500), a rabbit anti-pRab8A phospho T72 monoclonal antibody (Abcam 230260, 1:1000), and a rabbit anti-GAPDH monoclonal antibody (Cell Signalling Technology 14C10, 1:3000), diluted in 1% milk, overnight at 4ºC. The next day, the membranes were washed once with TBS-T for 10 minutes and three times at room temperature with TBS-T before the secondary antibodies were added. A secondary goat anti-rabbit Licor680 antibody (LI-COR biosciences) was diluted 1:5000 in 4% BSA and added to the top LRRK2-blotted membrane and incubated at room temperature for 1 hour. Secondary goat anti-rabbit Licor680 and goat anti-mouse Licor800 were diluted 1:5000 in 4% milk and incubated with the lower half of the membrane for 1 hour at room temperature. Membranes were washed for 10 minutes two times with TBS-T and once with TBS, and then imaged on an Odyssey CLx Li-Cor imaging system. After imaging, the lower half of the membrane was blocked again for 1 hour at room temperature with 5% milk and then incubated with primary rabbit anti-FLAG polyclonal antibody (ptg labs 20543-1-AP) diluted 1:500 in 1% milk overnight at 4ºC. The membrane was washed as before, then incubated at room temperature with secondary goat anti-rabbit Licor680 diluted 1:5000 in 4% milk. After washing again in TBS-T, the membrane was reimaged on the Li-Cor imaging system. Images were processed with ImageJ (version 2.14.0). The difference in phosphorylation of Rab8A between DMSO, MLi-2, and DARPin E11 conditions was analyzed using a one-way ANOVA and corrected using Tukey’s multiple comparison test. All statistical analyses were performed using GraphPad Prism (version 10.1.0).

## Results

### Identification of LRRK2-specific DARPins

To identify LRRK2-specific DARPins, we started with a library of N2C and N3C DARPins with randomized loops and capping repeats. For LRRK2, we used the well-characterized C-terminal catalytic half of LRRK2 that contains its ROC GTPase, COR-A and COR-B, Kinase, and WD40 domains that we refer to as LRRK2^RCKW^ (**Figure 1A**) (Deniston et al., 2020). We then used ribosome display with immobilized LRRK2^RCKW^ protein to enrich the DARPin pool for those that bound LRRK2 (Dreier and Plückthun, 2012). After four rounds of selection, 380 of the obtained DARPins were expressed in *E. coli* and probed in a homogeneous time-resolved fluorescence (HTRF) assay. Next, we scaled up our purification of the 20 top scoring hits (amino acid sequences are shown in **Table S1**) and performed size exclusion chromatography (SEC) (**Figure S1**) and Surface Plasmon Resonance (SPR) experiments to determine which DARPins to pursue further. One of these DARPins, E11, showed robust SPR data and was one of the highest affinity LRRK2 binders identified. DARPin E11 also interacted with one of LRRK2’s dimerization interfaces (see below), suggesting that it could provide a protein-based high-affinity tool for studying LRRK2 dimerization and filament formation in cells.

### Biophysical characterization of the LRRK2^RCKW^:E11 complex

To characterize DARPin E11, we first performed SPR experiments, applying DARPin E11 solutions at multiple concentrations to obtain association and dissociation kinetics (**Figure 1B, C**). The plateau values were fitted to a Langmuir model, and the K_D_ value was calculated to be 70 nM. We also probed the LRRK2^RCKW^:E11 interaction by bio-layer interferometry (BLI). Biotinylated LRRK2^RCKW^ was immobilized on the biosensors, and multiple concentrations of DARPin E11 were used. The dose-response curve was fitted to a Langmuir model, resulting in a K_D_ value of 11 nM (**Figure S2**). Together, our data indicate that DARPin E11 bound tightly to LRRK2^RCKW^ with K_D_ values in the low nanomolar range. The dissociation off-rate of the LRRK2^RCKW^:E11 complex (k_off_ in SPR ∼0.005 s^-1^) suggested that the complex is long-lived; this was supported by its stability in SEC experiments (**Figure 1D**). Finally, we compared LRRK2^RCKW^ alone to the LRRK2^RCKW^:E11 complex using mass photometry, where samples are at equilibrium. As expected, LRRK2^RCKW^ alone was mainly monomeric with a minor portion of dimers. The addition of a 2.5-fold excess DARPin E11 shifted the entire histogram to higher masses to a mass that corresponded to the addition of a DARPin molecule (**Figure 1E**). These results demonstrate the reversible formation of stable complexes with low nanomolar affinity between LRRK2^RCKW^ and DARPin E11.

### Cryo-EM structure of LRRK2^RCKW^ bound to DARPin E11

Next, we wanted to determine the binding mode between DARPin E11 and LRRK2. To do this, we prepared samples for analysis by cryo-EM by mixing LRRK2^RCKW^, GDP/Mg^2+^, a co-factor of LRRK2’s small G-protein binding (ROC) domain and DARPin E11. The workflow for data collection and analysis is summarized in **Figure S3**. The resulting cryo-EM map and model showed that DARPin E11 bound to the LRRK2 WD40 domain. The binding interface between DARPin E11 and LRRK2^RCKW^ was located at the bottom of the WD40 domain, adjacent to its central binding cavity and opposite the kinase active site (**Figure 2A**). Two insertions connecting neighboring blades of the WD40 ß-propeller (containing 7 blades) were particularly prominent in the binding interface: the helix-loop-helix motif connecting blades 4 and 5 (residues 2339 to 2351) and the partially disordered insertion connecting blades 5 and 6 (residues 2388 to 2414). The area of the binding interface was calculated using the PISA server resulting in an interaction interface of 877 A^2^, a value that is similar to what was calculated for other DARPin complexes (Blanc et al., 2023). In addition to hydrophobic interactions of the extensive binding interface, several key residues in LRRK2^RCKW^ and DARPin E11 were identified that formed polar interactions. (i) The backbone carbonyls from LRRK2^RCKW^ residues Gln2342, Leu2343 and Ser2345 formed a well-defined interaction network with residue Arg25 from DARPin E11 (**Figure 2B**). (ii) Several DARPin E11 residues including Val50, His54, Asp58, Asp79, Tyr81 and Leu88 formed a deep pocket that accommodated the side chain of the LRRK2^RCKW^ residue Tyr2346. The Asp79 side chain resided in the back of this pocket with the distance between Asp79 and the Tyr2346 hydroxy group allowing electrostatic interactions. Furthermore, sandwich π-π stacking between Tyr81 and Tyr2346 was observed, highlighting the importance of residue Tyr2346 in DARPin E11 binding (**Figure 2C**). (iii) Another key residue for the interaction was LRRK2^RCKW^ Arg2413. Its guanidino group was close to the DARPin E11 residues Phe83, Asp112 and His113, the latter again allowing for electrostatic interactions (**Figure 2D**). As expected, on the DARPin E11 side most interacting residues were situated in the randomized region of the core helices. Interestingly, several interacting residues including Arg25 and Asp58 were situated in the core helices (**Figure 2D**).

**Figure 2.**
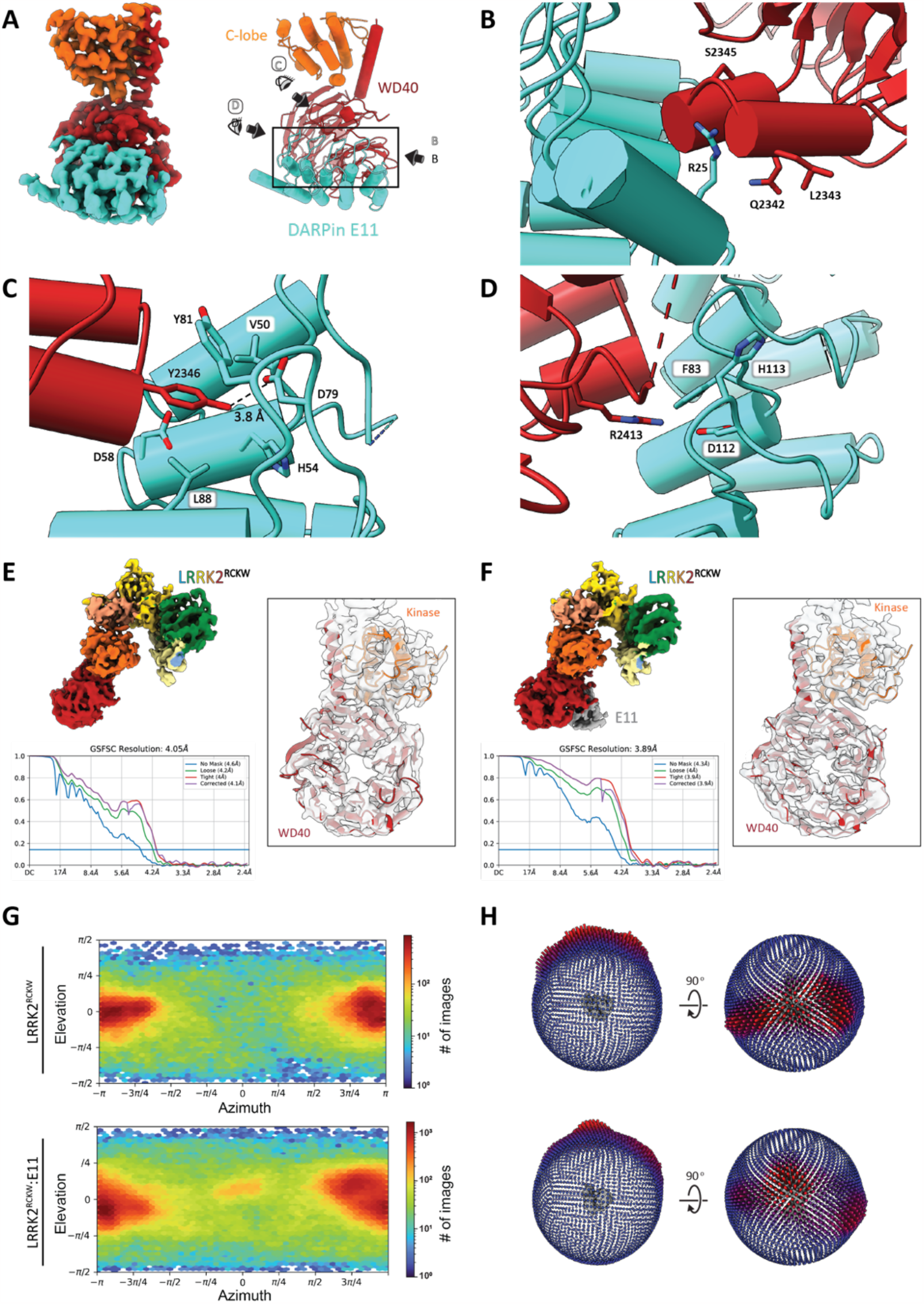
Cryo-EM structure of the LRRK2^RCKW^:DARPin E11 complex. **A**. Cryo-EM map and model of LRRK2^RCKW^ bound to DARPin E11. Because of the focused refinement strategy used to maximize the resolution of the DARPin-WD40 part of the structure, only the kinase C-lobe and the WD40 domain are seen in this map. E11 binds to the top of the WD40 domain opposite the kinase active site, next to the central WD40 pore. The eyes and arrows surrounding the rectangle on the right indicate the direction of the views shown in panels (B-D). **B-D**. Close-up views of the binding interface. Key residues for the interaction are highlighted. **E, F**. Cryo-EM maps, FSC plots, and model-to-map fits for LRRK2^RCKW^ alone or bound to DARPin E11. **G, H**. Plots of Euler angle distribution generated in cryoSPARC (left) or Relion (right) for LRRK2^RCKW^ alone (top) or bound to DARPin E11 (bottom). Addition of E11 increased the number of orientations adopted by LRRK2^RCKW^ on the cryo-EM grids.

### DARPin E11 reduces preferred orientation on cryo-EM grids

The DARPin E11 proved to be useful in a way we had not anticipated. When we determined the LRRK2^RCKW^:E11 structure, we noticed that the cryo-EM map of the complex showed better density for LRRK2^RCKW^ than what we had observed in the map of LRRK2^RCKW^ solved in the absence of E11 (**Figure 2E, F**). The reason for this improvement appeared to be a favorable random distribution of particle views in the presence of DARPin E11 (**Figure 2G, H**), although we cannot rule out an effect from the additional mass and features contributed by the DARPin. Given that both LRRK2^FL^ and LRRK2^RCKW^ have shown strong preferred orientations in the past (Deniston et al., 2020; Myasnikov et al., 2021), DARPin E11, and LRRK2-specific DARPins in general, have the potential to significantly improve the LRRK2 structures that can be obtained using cryo-EM.

### DARPin E11 does not alter the kinase activity of LRRK2^RCKW^ in vitro

We next sought to understand the functional implications of the LRRK2:DARPin E11 interaction. A subset of Rab GTPases, which mark specific membrane compartments in cells, are known substrates of LRRK2 (Steger et al., 2017, 2016). Thus, we determined if DARPin E11 binding to LRRK2^RCKW^ impacted its kinase activity. To do this, we performed in vitro kinase assays with LRRK2^RCKW^ and its substrate Rab8A in the presence of varying concentrations of DARPin E11 (**Figure 3A**). The ratio between the non-phosphorylated Rab8A and pRab8A was determined using electrospray time-of-flight (tof) mass spectrometry. These assays revealed that even at the saturating concentration of 1 μM, DARPin E11 had no effect on LRRK2^RCKW^ kinase activity (**Figure 3B**).

**Figure 3.**
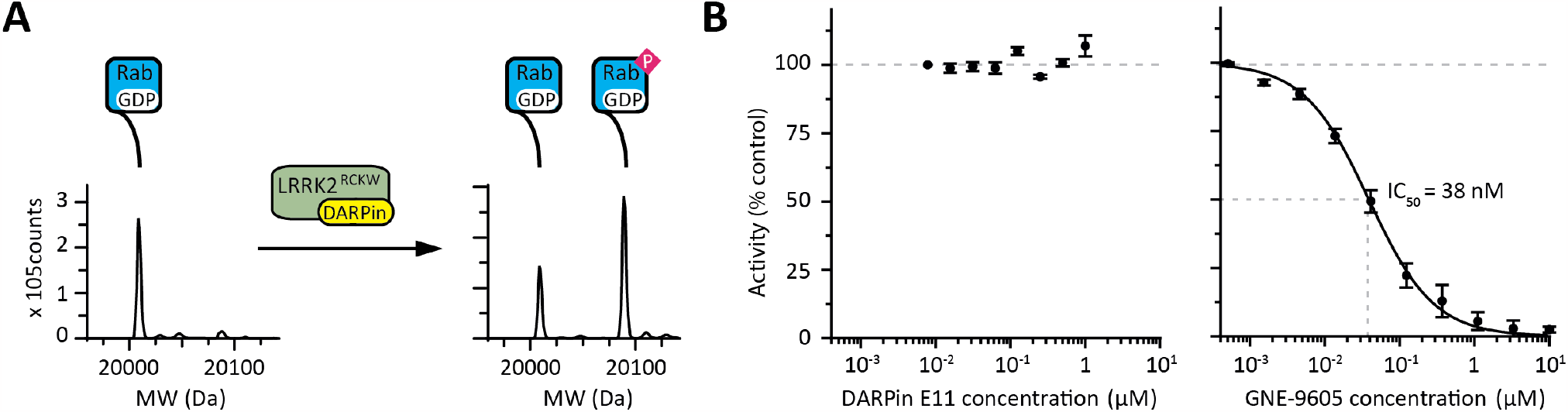
DARPin E11 does not affect the kinase activity of LRRK2^RCKW^ in vitro. **A**. In-vitro assay measuring the level of Rab8A phosphorylation by LRRK2^RCKW^. Rab8A and LRRK2^RCKW^ are incubated in the presence of ATP and kinase activity is monitored by measuring the levels of substrate (Rab8A) and product (pRab8A) by mass spectrometry. **B**. Kinase activity of LRRK2^RCKW^ in the presence of either DARPin E11 (left) or the LRRK2-specific kinase inhibitor GNE-9605 (right).

Full length LRRK2 (LRRK2^FL^) has been reported to adopt an autoinhibited conformation (Myasnikov et al., 2021). The mechanism of autoinhibition has been proposed to be mediated by the N-terminal LRR domain folding over the kinase active site, a structural arrangement that partially blocks the substrate binding site of LRRK2. Folding of the LRR domain over the kinase active site is stabilized by interactions with the WD40 domain and the C-terminal helix of LRRK2. Therefore, we hypothesized that DARPin E11 interacting with the WD40 domain might interfere with the autoinhibited conformation. However, the overlay of the autoinhibited LRRK2^FL^ model (Myasnikov et al., 2021) and our LRRK2^RCKW^:E11 model showed that this was not the case, as DARPin E11 could bind to the WD40 domain without introducing clashes with the LRR (**Figure 4A**). However, the DARPin E11 binding site on LRRK2^RCKW^ did overlap with the site of WD40:WD40 dimerization (**Figure 4B**).

**Figure 4.**
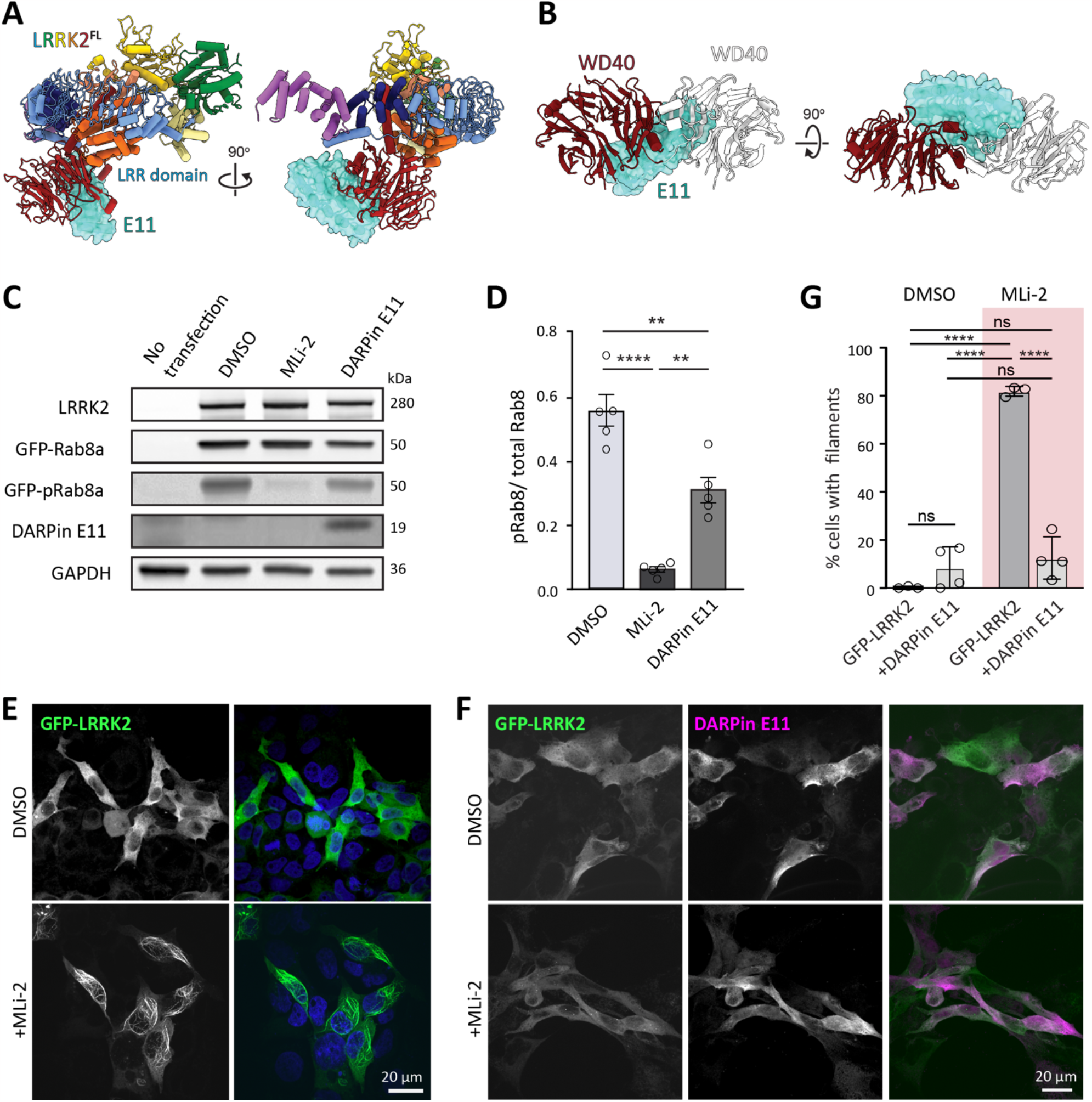
DARPin E11 disrupts LRRK2^FL^ filament formation and decreases Rab8 phosphorylation. **A**. Modeling of E11 bound to the autoinhibited conformation of LRRK2^FL^ shows no steric clashes between E11 and LRRK2. **B**. The binding site of DARPin E11 on the WD40 domain overlaps with the WD40:WD40 dimerization interface that is involved in the formation of microtubule associated LRRK2 filaments. **C**. Rab8 phosphorylation in 293T cells overexpressing LRRK2^FL^ and GFP-Rab8, with or without DARPin E11-3x FLAG. 293T cells were transiently co-transfected with LRRK2^FL^ and GFP-Rab8 or LRRK2^FL^, GFP-Rab8, and DARPin E11-3xFLAG for 48 hours. Cells transfected only with LRRK2^FL^ and GFP-Rab8 were treated with DMSO or 2 μM MLi-2 for 1 hour. Cells were lysed and immunoblotted for phospho-Rab8 (pT72), total GFP-Rab10, total LRRK2, DARPin E11-3xFLAG and GAPDH. **D**. Quantification from five western blots plotting the ratio of GFP-pRab8 to total GFP-Rab8. Statistics were generated in GraphPad using a one-way ANOVA analysis with a Tukey’s multiple comparison of means. **** p <0.0001 DMSO and MLi-2, ** p=0.0013 DMSO and DARPin E11, ** p= 0.0012 MLi-2 and DARPin E11. **E, F**. Representative images of 293T cells expressing either GFP-LRRK2^FL^ (C) or GFP-LRRK2^FL^ and DARPin E11 (D), treated with DMSO or MLi-2 for 2 hours. **G**. Quantification of the percent cells (mean +/-sd) with GFP-LRRK2^FL^ filaments in the presence or absence or DARPin E11. Each data point on the graph represents a replicate. 3 or 4 independent replicates were done per condition, with each replicate containing 48-140 cells. Statistics were generated using a One Way ANOVA with a Tuckey’s multiple comparison of means. **** p < 0.0001. ns, not significant.

### DARPin E11 decreases LRRK2 kinase activity in cells

Our experiments with LRRK2^RCKW^ showed that DARPin E11 did not affect LRRK2 kinase activity in vitro. Next, we wanted to determine if DARPin E11 affected LRRK2 kinase activity in cells. We transfected 293T cells with full-length untagged LRRK2 and GFP-Rab8A with or without co-transfection of DARPin E11-3xFLAG and monitored the phosphorylation of GFP-Rab8A using a phospho-specific Rab8A antibody. Strikingly, in contrast to what we observed in vitro using the N-terminally truncated LRRK2^RCKW^ construct, the presence of DARPin E11 significantly reduced LRRK2^FL^ kinase activity in cells compared to the DMSO control (**Figure 4C, D**).

### DARPin E11 disrupts LRRK2 microtubule association

LRRK2 interact with microtubules in cells under some circumstances (Kett et al., 2012). These LRRK2 filaments have been observed by cryo-electron tomography (cryo-ET) in cells for LRRK2^FL^ (Watanabe et al., 2020) and in vitro for LRRK2^RCKW^ (Snead et al., 2022). The tendency of LRRK2 to form filaments is increased by most LRRK2 PD mutations (Kett et al., 2012) and by the presence of type-I kinase inhibitors that stabilize the LRRK2 active conformation (Blanca Ramírez et al., 2017; Deniston et al., 2020; Kett et al., 2012; Schmidt et al., 2021). The LRRK2 filaments are formed by LRRK2 dimers, mediated by a COR-B:COR-B interface, which interacts through a WD40:WD40 interface (Deniston et al., 2020; Snead et al., 2022; Watanabe et al., 2020). This interface is best resolved in a crystal structure of WD40 dimers (Zhang et al., 2019). The overlay with our LRRK2^RCKW^:E11 complex showed that the presence of DARPin E11 was incompatible with the formation of the WD40:WD40 dimer (**Figure 4A**). Thus, we hypothesized that DARPin E11 would prevent the formation of LRRK2 filaments in cells. To test this, we co-expressed DARPin E11 and GFP-LRRK2^FL^ in 293T cells and treated the cells with the LRRK2-specific type-I inhibitor MLi-2, which as expected induced LRRK2 filament formation on microtubules (**Figure 4E, F**). While LRRK2 filaments formed on microtubules in over 80% of control cells transfected with GFP-LRRK2^FL^ and treated with MLi-2, cells co-expressing DARPin E11 did not form filaments in the presence of MLi-2, suggesting that DARPin E11 disrupts WD40-dependent LRRK2 dimerization and its association with microtubules (**Figure 4G**).

## Discussion

Here we described the development of LRRK2-specific DARPins. The development of these small protein-based probes for LRRK2 adds important reagents to the growing toolbox to explore LRRK2 function in health and disease. We report the sequence of 20 LRRK2 DARPins and characterize DARPin E11 as a protein-based affinity reagent that binds to the WD40 domain of LRRK2.

Unexpectedly, we found that binding of DARPin E11 to LRRK2’s WD40 domain decreases preferred orientations on cryo-EM grids, allowing us to solve a cryo-EM structure with improved resolution. Thus, DARPin E11, and potentially other LRRK2-specific DARPins, will facilitate future cryo-EM studies of LRRK2. Given the higher resolution obtained in the presence of DARPin E11, this may be particularly useful for solving structures of LRRK2 in complex with small molecules, such as kinase inhibitors.

Our studies of DARPin E11 revealed that this affinity reagent can be used to study intra- and inter-molecular interactions modulating LRRK2 functions in cells. For instance, we showed that DARPin E11 disrupted the ability of LRRK2 to form microtubule-associated filaments, which form in the presence of the LRRK2 kinase inhibitor MLi-2 and are enhanced by many PD mutations (Kett et al., 2012). Thus, DARPin E11 could be used as a tunable reagent in cells to prevent the formation of the WD40-WD40 interface. In the future this reagent could be used to study endogenous LRRK2 in PD relevant cell types.

In terms of LRRK2’s cellular function, our most intriguing observation was that even though DARPin E11 had no effect on the kinase activity of truncated LRRK2^RCKW^ in vitro, its expression in cells led to reduced phosphorylation of Rab8A, one of LRRK2’s physiological substrates. This suggests that DARPin E11 could be competing with accessory factors that interact with the WD40 domain to control LRRK2’s subcellular localization, orientation, and/or conformation. This idea is consistent with the properties of a rare LRRK2 variant (V2390M) identified in a male PD patient from Spain (Clarimón et al., 2008). This variant showed reduced phosphorylation of Rab10 in cells (Kalogeropulou et al., 2022). While the small valine side chain (V2390) is accommodated well in our structure of the LRRK2^RCKW^:E11 complex, the bulkier methionine side chain has the potential to disrupt, or at least weaken the binding of E11 to LRRK2’s WD40 domain. We speculate that DARPin E11 and the V2390M variant may be disrupting the same docking site for cellular LRRK2 regulators. The identity of these regulators and how they modulate the LRRK2 kinase activity remains to be discovered, but DARPin E11 is a tool that could lead to their discovery.

The observations summarized above also suggest a new approach to design small-molecule inhibitors of LRRK2’s kinase activity in cells by targeting its WD40 domain rather than its kinase: small-molecule libraries could be screened for binders that disrupt the interaction between LRRK2’s WD40 domain and DARPin E11. The identified small-molecule binders would serve as starting points for the development of high-affinity binders of the LRRK2 WD40 domain that may be potent inhibitors of LRRK2 in cells and could be further developed as therapeutics for PD.

## Acknowledgments

The authors are grateful to Sven Furler, Thomas Reinberg und Joana Marinho for carrying out the ribosome display selections and DARPin screening. This research was funded in whole or in part by Aligning Science Across Parkinson’s (ASAP-000519) through the Michael J. Fox Foundation for Parkinson’s Research (MJFF). SK is grateful for support by the SGC, a registered charity (no: 1097737) that receives funds from AbbVie, Bayer AG, Boehringer Ingelheim, Canada Foundation for Innovation, Eshelman Institute for Innovation, Genentech, Genome Canada, EU/EFPIA/OICR/McGill/ KTH/Diamond, Innovative Medicines Initiative 2 Joint Undertaking (EUbOPEN grant 875510), Janssen, Pfizer and Takeda. SLRP is also supported by the Howard Hughes Medical Institute.

## Author contributions

VD, DC, FP, and KRAA conducted biochemical studies and biophysical assays. MSM solved and refined cryo-EM structures; CG and WV profiled DARPins using BLI; SLRP, EPK, KSH, and LVN conducted cellular imaging and kinase studies; BD and AP designed and supervised the selection and screening of DARPins; SM, AP, SLRP, AEL, and SK supervised the research. SM, SK, AEL, and SLRP wrote the manuscript, which was edited and approved by all authors.

## Data availability

The cryo-EM map and model for LRRK2^RCKW^:E11 have been deposited in the EM and Protein Data Banks, respectively. Accession codes are 8U1B (PDB) and EMBD-41806 (EMDB).

