## Supplemental Material for "Development of LRRK2 designed ankyrin-repeat proteins"

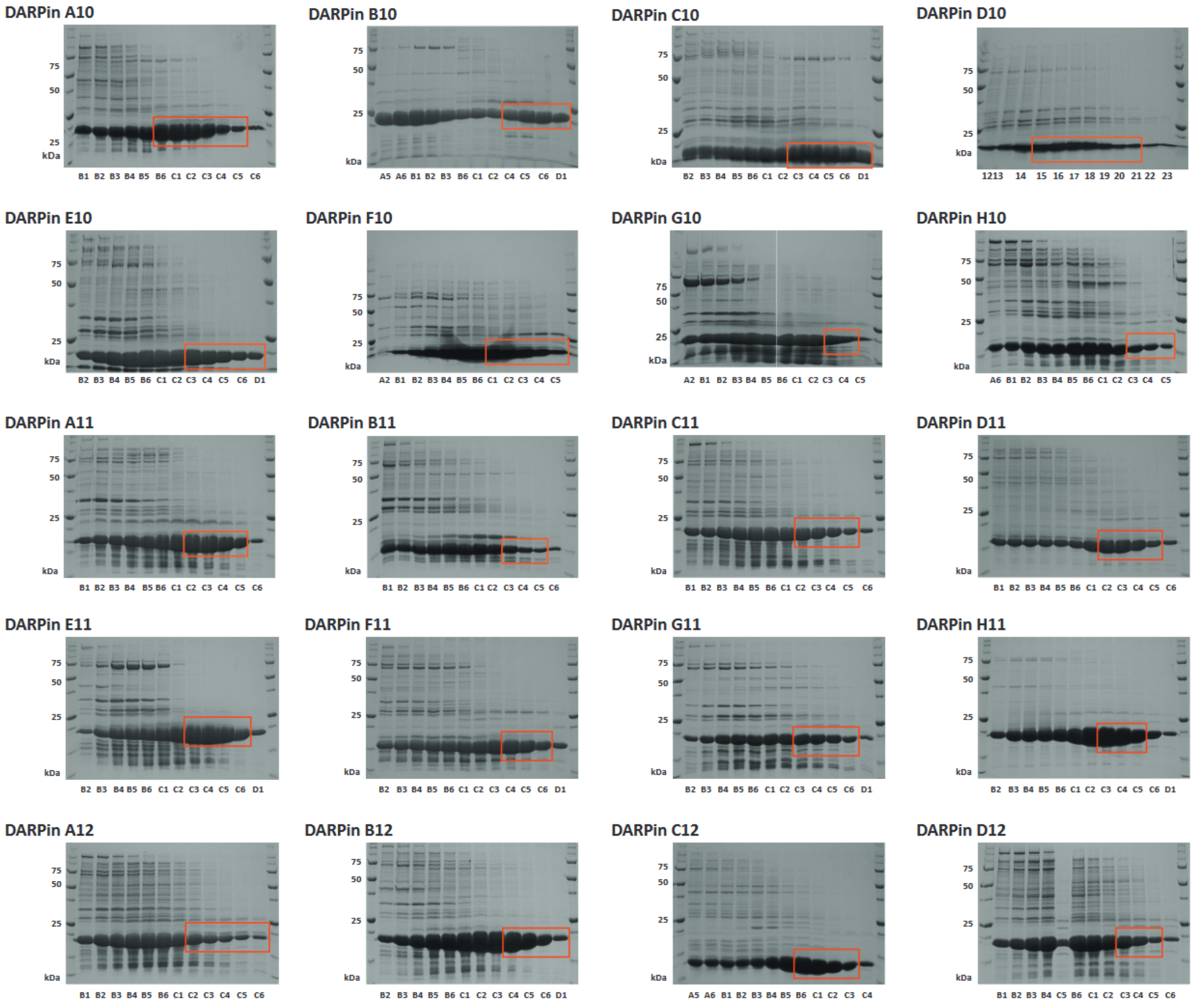

**Figure S1. Analysis of DARPins by size-exclusion chromatography (SEC).**
SEC for the 20 top DARPins from our screen. The eluting fractions were collected (3 mL samples) and analyzed by SDS PAGE. The red boxes indicate the fractions that were pooled and stored for follow-up experiments.

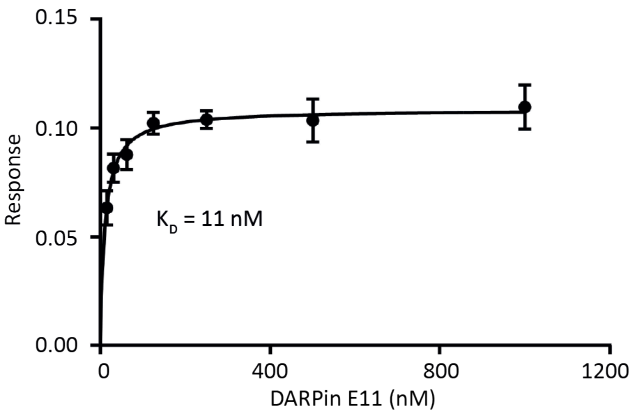

**Figure S2. DARPin E11 binds to LRRK2$^{RCKW}$ with high affinity.**
Bio-layer interferometry analysis of the LRRK2$^{RCKW}$:E11 interaction yielded a $K_D$ value of 11 nM. The Langmuir equation was used to fit the dose-response curve.

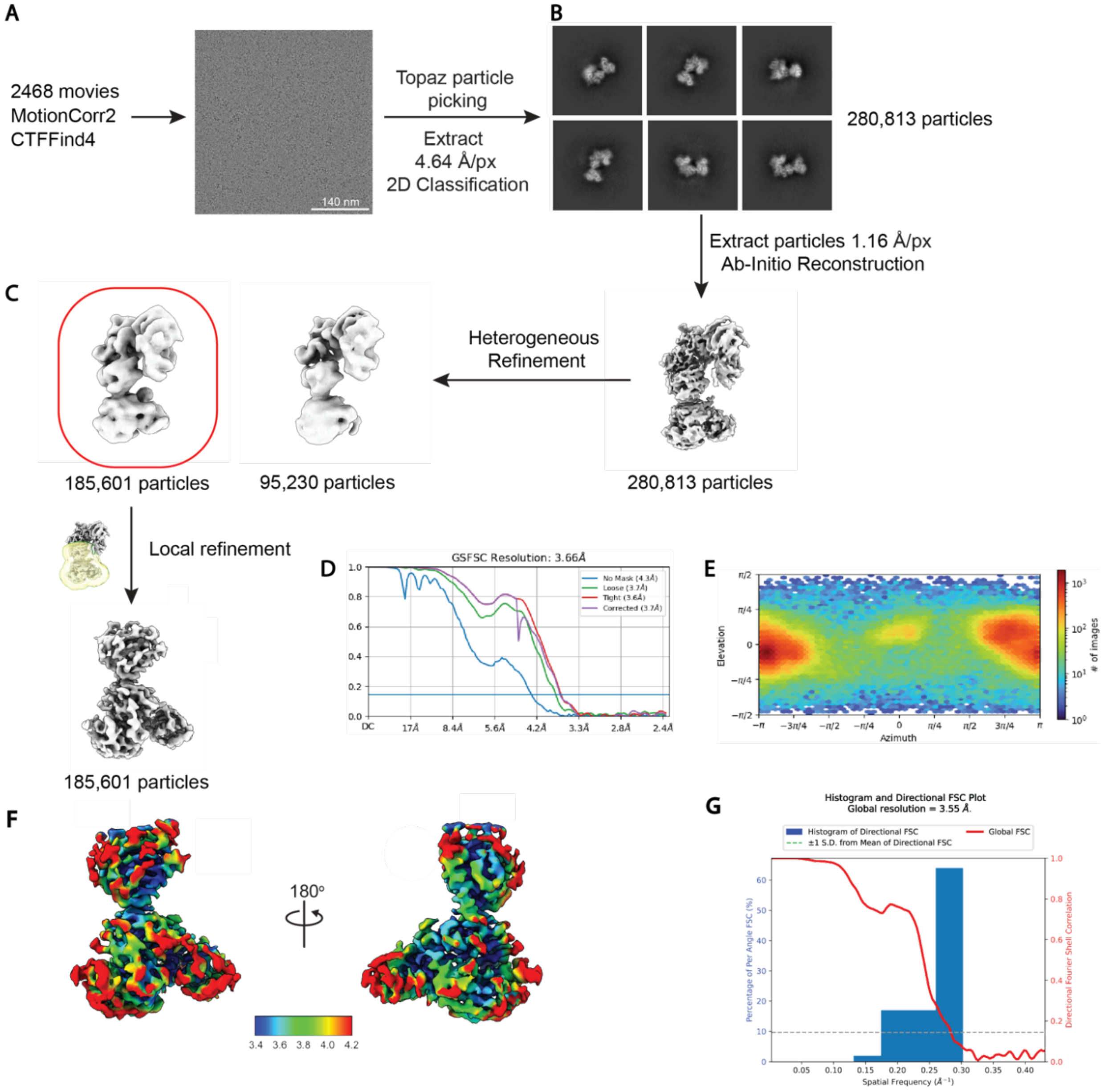

**Figure S3. Cryo-EM data processing workflow for LRRK2$^{RCKW}$:E11 DARPin.**
**A.** An initial dataset with 2,468 movies was collected from a sample of the LRRK2$^{RCKW}$:E11 DARPin complex. A typical micrograph is shown. **B.** 2D class averages. **C.** Data processing strategy. All processing was done in cryoSPARC. **D.** FSC curves. **E.** Euler angle distribution. **F.** Local Resolution map. **G.** 3D-FSC analysis including global half map FSC (red line) and histogram of values evenly sampled over the 3D FSC (blue bars).

**Table S1. DARPin amino acid sequences of 20 DARPins selected for scale-up purification.**

| DARPin | Sequence |
|---|---|
| A10 | MRGSHHHHHHHHGSDLGKKLLEAARAGQDDEVRILMANGADVNAWDWMGKTPLHLAAHDGHLEIVEVLLKTG<br>ADVNARDTIGITPLHLTAWKGHLEIVEVLLKHGADVNAADWYGMTPLHLAAYEGHLEIVEVLLKHGADVNAQ<br>DWTGSTPFDLAAYRGNEDIAEVLQKAAKLNDYKDDDDK |
| B10 | MRGSHHHHHHHHGSDLGKKLLEAARAGQDDEVRILMANGADVNAVDWSGYTPLHLAALAGHLEIVEVLLKTG<br>ADVNAEDTPLHLAADEGHLEIVEVLLKTGADVNADDSYGVTPLHLAAWNGHLEIVEVLLKAGADVNATDNQG<br>RTPLHLAAMHGNEDIAEVLQKAAKLNDYKDDDDK |
| C10 | MRGSHHHHHHHHGSDLGKKLLEAAMKGQDDEVRILMANGADVNAYDAWGYTPLHLAAYTGHLEIVEVLLKTG<br>ADVNATDERGVTPLHLAAWLGHLEIVEVLLKTGADVNAQDVWGTTPLHLAAQAGHLEIVEVLLKHGADVNAQ<br>DFTGWTPFDLAASVGNEDIAEVLQKAAKLNDYKDDDDK |
| D10 | MRGSHHHHHHHHGSDLGKKLLEAAVYGQDDEVRILMANGADVNAVDMNGETPLHLATIQGHLEIVEVLLKTG<br>ADVNAEDWIGATPLHLAAIWGHLEIVEVLLKHGADVNALDISGATPFDLAAMIGNEDIAEVLQKAAKLNDYK<br>DDDDK |
| E10 | MRGSHHHHHHHHGSDLGKKLLEAAKEGQDDEVRILMANGADVNAWDKFGMTPLHLAAVSGHLEIVEVLLKTG<br>ADVNASDYEGDTPLHLAAAWGHLEIVEVLLKAGTDVNASDYWGYTPLHLAAWAGHLEIVEVLLKHGADVNAQ<br>DWQGSTPFDLAASSGNEDIAEVLQKAAKLNDYKDDDDK |
| F10 | MRGSHHHHHHHHGSDLGKKLLEAAWMGQLDEVRILMANGADVNAEDMFGSTPLHLAANAGHLEIVEVLLKAG<br>ADVNAEDRFGITPLHLAAHVGHLEIVEVLLKHGADVNAQDSFGWTPFDLAAWHGNEDIAEVLQKAAKLNDYK<br>DDDDK |
| G10 | MRGSHHHHHHHHGSDLGKKLLEAAAYGQDDEVRILMANGADVNATDQWGFTPLHLAAFNGHLEIVEVLLKTG<br>ADVNADDVVGQTPLHLAAWQGHLEIVEVLLKAGADVNAWDFWGQTPLHLAAQLGHLEIVEVLLKHGADVNAQ<br>DDWGETPFDLAIDNGNEDIAEVLQKAAKLNDYKDDDDK |
| H10 | MRGSHHHHHHHHGSDLGKKLLEAARAGQDDEVRILMANGADVNAADWIGRTPLHLAAMHGHLEIVEVLLKTG<br>ADVNAFDWYGVTPLHLAAYNGHLEIVEVLLKTGADVNAQDYFGITPLHLAAYWGHLEIVEVLLKHGADVNAQ<br>DIAGVTPFDLAIDNGNEDIAEVLQKAAKLNDYKDDDDK |
| A11 | MRGSHHHHHHHHGSDLGKKLLEAARAGQDDEVRILMANGADVNAVDMWGSTPLHLAATDGHLEIVEVLLKTG<br>ADVNAWDLMGLTPLHLAAVWGHLEIVEVLLKAGADVNAWDMWGWTPLHLAADQGHLEIVEVLLKHGADVNAQ<br>DKFGKTPFDLAIDNGNEDIAEVLQKAAKLNDYKDDDDK |
| B11 | MRGSHHHHHHHHGSDLGKKLLEAAVTGQDDEVRILMANGADVNAADEWGDTPLHLAANYGHLEIVEVLLKHG<br>ADVNAQDYFGITPLHLAAVIGHLEIVEVLLKHGADVNAQDWAGWTPFDLAAWTGNEDIAEVLQKAAKLNDYK<br>DDDDK |
| C11 | MRGSHHHHHHHHGSDLGKKLLEAARAGQDDEVRILMANGADVNADDNNGVTPLHLAADHGHLEIVEVLLKTG<br>ADVNAMDIYGLTPLHLAAWLGHLEIVEVLLKTGADVNAVDNTGWTPLHLAAFLGHLEIVEVLLKHGADVNAQ<br>DFWGFTPFDLAAITGNEDIAEVLQKAAKLNDYKDDDDK |
| D11 | MRGSHHHHHHHHGSDLGKKLLEAARAGQDDEVRILMANGADVNASDWWGHTPLHLAAAKGHLEIVEVLLKTG<br>ADVNAVDIRGVTPLHLAAWQGHLEIVEVLLKAGADVNAQDVWGWTPLHLAAIWGHLEIVEVLLKHGADVNAQ<br>DKFGKTPFDLAIDNGNEDIAEVLQKAAKLNDYKDDDDK |
| E11 | MRGSHHHHHHHHGSDLGKKLLEAARAGQDDEVRILMANGADVNATDEAGVTPLHLAADSGHLEIVEVLLKTG<br>ADVNAWDHYGFTPLHLAAHVGHLEIVEVLLKAGADVNAQDHAGWTPLHLAALYGHLEIVEVLLKHGADVNAQ<br>DMWGETPFDLAIDNGNEDIAEVLQKAAKLNDYKDDDDK |
| F11 | MRGSHHHHHHHHGSDLGKKLLEAAWIGQDDEVRILMANGADVNAVDWWGITPLHLAASSGHLEIVEVLLKTG<br>ADVNADDHWGSTPLHLAAWFGHLEIVEVLLKAGADVNAYDEVGHTPLHLAIDNGNEDIAEVLQKAAKLNDYK<br>DDDDK |
| G11 | MRGSHHHHHHHHGSDLGKKLLEAARAGQDDEVRILMANGADVNAMDVVGMTPLHLAAVSGHLEIVEVLLKTG<br>ADVNATDWWGYTPLHLAASFGHLEIVEVLLKAGADVNARDQLGDTPLHLAANVGHLEIVEVLLKHGADVNAQ<br>DMYGDTPFDLAAWVGNEDIAEVLQKAAKLNDYKDDDDK |
| H11 | MRGSHHHHHHHHGSDLGKKLLEAARAGQDDEVRILMANGADVNAVDWWGVTPLHLAANEGHLEIVEVLLKTG<br>ADVNAYDNLGATPLHLAAAKGHLEIVEVLLKTGADVNAWDYWGWTPLHLAAMRGHLEIVEVLLKHGADVNAQ<br>DLHGTTPFDLAAHNGNEDIAEVLQKAAKLNDYKDDDDK |
| A12 | MRGSHHHHHHHHGSDLGKKLLEAAVHGHLDEVRILMANGADVNAEDEHGWTPLHLAAAYGHLEIVEVLLKTG<br>ADVNAWDWDGHTPLHLAAYIGHLEIVEVLLKTGADVNAQDAWGWTPLHLAAVQGHLEIVEVLLKHGADVNAY<br>DHFGQTPFDLAAWVGNEDIAEVLQKAAKLNDYKDDDDK |
| B12 | MRGSHHHHHHHHGSDLGKKLLEAARAGQDDEVRILMANGADVNAVDRWGITPLHLAAVTGHLEIVEVLLKTG<br>ADVNATDWWGWTPLHLAAMEGHLEIVEVLLKAGADVNAWDDWGDTPLHLAAITGHLEIVEVLLKHGADVNAQ<br>DKFGKTPFDLAIDNGNEDIAEVLQKAAKLNDYKDDDDK |
| C12 | MRGSHHHHHHHHGSDLGKKLLEAAMSGQDDEVRILMANGADVNTQDQFGLTPLHLAAIEGHLEIVEVLLKTG<br>ADVNALDWVGTTPLHLAAYVGHLEIVEVLLKHGADVNEDIAEVLQKAAKLNDYKDDDDK |
| D12 | MRGSHHHHHHHHGSDLGKKLLEAARAGQDDEVRILMANGADVNALDFSGQTPLHLAAQWGHLEIVEVLLKTG<br>ADVNAADAWGTTPLHLAAYSGHLEIVEVLLKAGADVNAMDWWGITPLHLAAILGHLEIVEVLLKHGADVNAQ<br>DWQGFTPFDLAAWVGNEDIAEVLQKAAKLNDYKDDDDK |

**Table S2. Cryo-EM data collection, refinement and validation statistics.**

| | LRRK2$^{KW}$:DARPin E11 (EMDB-41806, PDB 8U1B) |
|---|---|
| **Data collection and processing** | |
| Magnification | 36000 |
| Voltage (kV) | 200 |
| Electron exposure (e–/Å$^2$) | 52 |
| Defocus range (μm) | -1.0 to -2.5 |
| Pixel size (Å) | 1.16 |
| Symmetry imposed | C1 |
| Initial particle images (no.) | 312,696 |
| Final particle images (no.) | 185,601 |
| Map resolution (Å) | 3.66 |
| FSC threshold | 0.143 |
| Map resolution range (Å) | 3.4 – 4.2 |
| **Refinement** | |
| Initial model used (PDB code) | 6VP7 |
| Model resolution (Å) | 3.66 |
| FSC threshold | 0.143 |
| Model composition | |
| Non-hydrogen atoms | 4540 |
| Protein residues | 604 |
| *B* factors (Å$^2$) | |
| Protein | 778.59 |
| R.m.s. deviations | |
| Bond lengths (Å) | 0.009 |
| Bond angles (°) | 1.338 |
| Validation | |
| MolProbity score | 2.34 |
| Clashscore | 18.15 |
| Ramachandran plot | |
| Favored (%) | 91.29 |
| Allowed (%) | 8.36 |
| Disallowed (%) | 0.35 |
